# Design and Validation of the First-in-Class PROTACs for Targeted Degradation of the Immune Checkpoint LAG-3

**DOI:** 10.1101/2025.08.09.669498

**Authors:** Nelson García Vázquez, Somaya A. Abdel-Rahman, Hossam Nada, Moustafa Gabr

## Abstract

Lymphocyte activation gene-3 (LAG-3) is an inhibitory immune checkpoint receptor that plays a central role in T cell exhaustion and immune evasion in cancer. While monoclonal antibodies targeting LAG-3 have entered clinical development, small molecule approaches remain largely unexplored. Here, we report the design and validation of the first-in-class PROTACs for targeted degradation of LAG-3. In this study, we repurposed a LAG-3-binding small molecule identified through DNA-encoded library (DEL) screening as the targeting ligand for a series of CRL4^CRBN^-based PROTACs designed with varied linker lengths. Western blot analysis in Raji-LAG3 cells demonstrated that **LAG-3 PROTAC-1** and **LAG-3 PROTAC-3** induce potent, dose-dependent degradation of LAG-3, with DC_50_ values of 274 nM and 421 nM, respectively. Molecular docking and molecular dynamics (MD) simulations revealed the LAG-3 binding mode of designed PROTACs and provided structural insights into PROTAC-mediated ternary complex formation. Collectively, this work establishes a proof-of-concept for chemical degradation of LAG-3 for the first time and paves the way for novel immunotherapeutic strategies.

---

Immune checkpoint inhibition has revolutionized cancer immunotherapy, delivering remarkable clinical outcomes and prolonged survival for select patient groups.^1^ Central to this progress are monoclonal antibodies (mAbs) that disrupt inhibitory signaling pathways mediated by checkpoint proteins, especially PD-1 and CTLA-4.^2^ Nevertheless, a significant proportion of patients either fail to respond or eventually relapse due to acquired resistance, posing a major limitation to the long-term success of ICB.^3^ To overcome these therapeutic barriers, recent efforts have turned toward the identification and targeting of alternative immune checkpoints to widen the spectrum of patients who may benefit from immunotherapy.^1^

Lymphocyte activation gene-3 (LAG-3) functions as a co-inhibitory receptor predominantly found on dysfunctional or exhausted T cells, making it an attractive immunotherapeutic target.^4–6^ It operates synergistically with PD-1 and CTLA-4 to amplify regulatory T cell–mediated suppression and dampen immune responses via antigen-presenting cells (APCs).^7^ LAG-3 binds to ligands including MHC class II molecules and the soluble protein FGL1, attenuating TCR signaling and thereby reducing T cell activation and cytokine production.^4^ Notably, FGL1 has been recognized as a non-classical ligand of LAG-3, contributing to immune evasion through suppression of antigen-specific T cell responses.^8^

Elevated LAG-3 expression has been documented in tumor-infiltrating lymphocytes (TILs) across multiple cancer types.^9,10^ Dual blockade strategies targeting PD-1 and LAG-3 have demonstrated synergistic enhancement of antitumor responses in preclinical models, including ovarian, colorectal, and melanoma tumors.^11^ Remarkably, the combination therapy of nivolumab (anti–PD-1) and relatlimab (anti–LAG-3) earned FDA approval in 2022 for use in advanced melanoma.^12^

Therapeutic targeting of LAG-3 in ongoing clinical trials remains predominantly limited to mAbs.^13^ While mAbs provide high specificity, their use is constrained by challenges such as suboptimal tissue distribution, elevated production costs, and the potential for immune-related adverse effects due to prolonged systemic exposure.^14,15^ In contrast, small molecule therapeutics offer distinct advantages, including oral administration, deeper tumor penetration, and tunable pharmacokinetic properties.^16^ These features may support more flexible dosing strategies and potentially reduce the incidence or severity of immune-related adverse effects. Moreover, small molecules can be designed to engage intracellular targets, offering opportunities for novel mechanisms of immune modulation. Thus, the development of small molecule LAG-3 modulators, particularly those capable of degrading the protein, could open new avenues in cancer immunotherapy, both as monotherapies and in rational combination strategies.

Proteolysis-targeting chimeras (PROTACs) represent a therapeutic modality that leverages the ubiquitin–proteasome system (UPS) to eliminate, rather than inhibit, disease-relevant proteins.^17–19^ A PROTAC molecule is composed of three modular components: a ligand that binds the protein of interest (POI), a ligand that recruits an E3 ubiquitin ligase, and a chemical linker that bridges the two.^17^ Upon binding both the POI and E3 ligase, the PROTAC induces formation of a ternary complex, facilitating ubiquitination of the target protein and its subsequent degradation by the proteasome. Importantly, PROTACs act catalytically and can be recycled following target degradation.^17^

This approach has gained remarkable traction as a rational drug discovery strategy, enabling the pharmacological removal of proteins previously considered undruggable.^18,19^ While PROTACs have shown promise in oncology, inflammation, and neurodegeneration, their application to immuno-oncology targets, particularly immune checkpoints, remains nascent. Given the therapeutic relevance of LAG-3 and other immune suppressive receptors, targeted degradation offers a unique opportunity to overcome the limitations of antibody-based approaches and expand the immunomodulatory toolkit. We recently employed DNA-encoded library (DEL) screening to identify compound **1** (Figure 1) as a LAG-3-binding hit.^20^ Subsequent structure-based optimization led to the development of compound **2** (Figure 2), a lead compound with the ability to block LAG-3 interactions in cellular and co-culture assays.^20^ This work paved the way for future research focused on targeting LAG-3 with small molecules to develop the next generation cancer immunotherapies.

**Figure 1.**
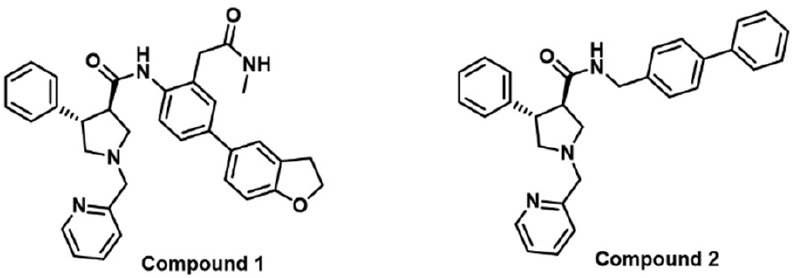
Chemical structures of compounds **1** and **2**.

**Figure 2.**
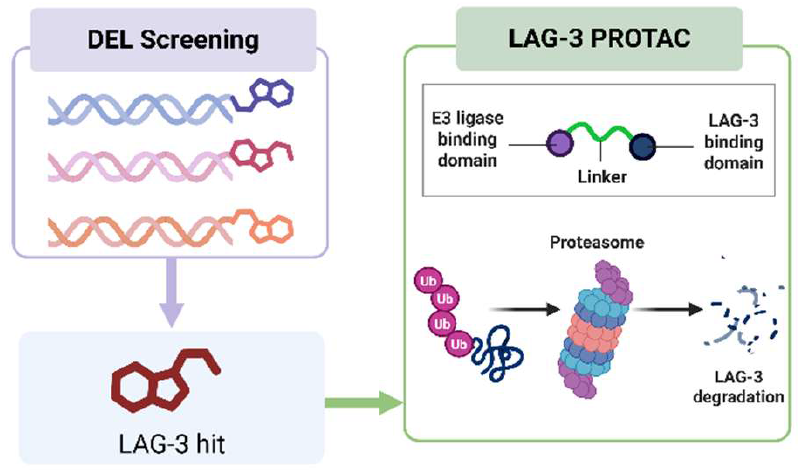
LAG-3 small molecule binder from DEL screening was developed into a PROTAC to induce LAG-3 degradation.

The identification of a DEL-derived LAG-3 binder (compound **1**) and its subsequent optimization into compound **2** provided a strong rationale for repurposing this chemical scaffold as a ligand for targeted protein degradation. In the present study, we build on this foundation by employing this scaffold as the LAG-3-binding moiety in the rational design of PROTACs. By linking it to E3 ligase recruiters through various linkers, we aim to generate bifunctional molecules capable of inducing LAG-3 ubiquitination and degradation via the proteasome (Figure 2). To our knowledge, this work represents the first report of PROTACs designed to degrade LAG-3, establishing a new direction for immunomodulation through targeted LAG-3 degradation.

Based on this strategy, we designed and synthesized LAG-3-targeted PROTACs by conjugating the pyrrolidine-3-carboxamide core (from compounds **1** and **2**) to pomalidomide as the E3 ligase recruiter for CRL4^CRBN^-mediated ubiquitination and degradation. The synthesis proceeded by amide coupling of the pyrrolidine-3-carboxylic acid derivative (Fragment **I**) and corresponding amine (Fragment **II**) using HBTU (Hexafluorophosphate Benzotriazole Tetramethyl Uronium) in DMF and in the presence of triethylamine (Scheme 1).

**Scheme 1.**
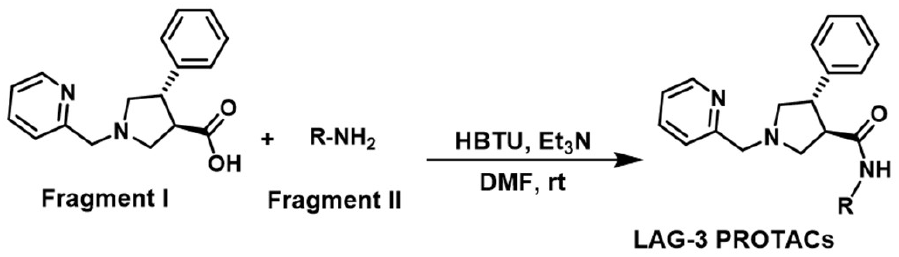
Synthesis of the designed LAG-3 PROTACs based on amide coupling of Fragments I and II.

The chemical structures of the synthesized LAG-3 PROTACs (**LAG-3 PROTAC-1, LAG-3 PROTAC-2**, and **LAG-3 PROTAC-3**) are shown in Figure 3. These molecules were designed with varied linker lengths to systematically assess how subtle changes in linker architecture influence degradation outcomes. The design rationale centered on preserving the LAG-3 scaffold while enabling productive ternary complex formation and ubiquitin transfer. The structural characterization of the synthesized PROTAC molecules is depicted in Figures S1-S8.

**Figure 3.**
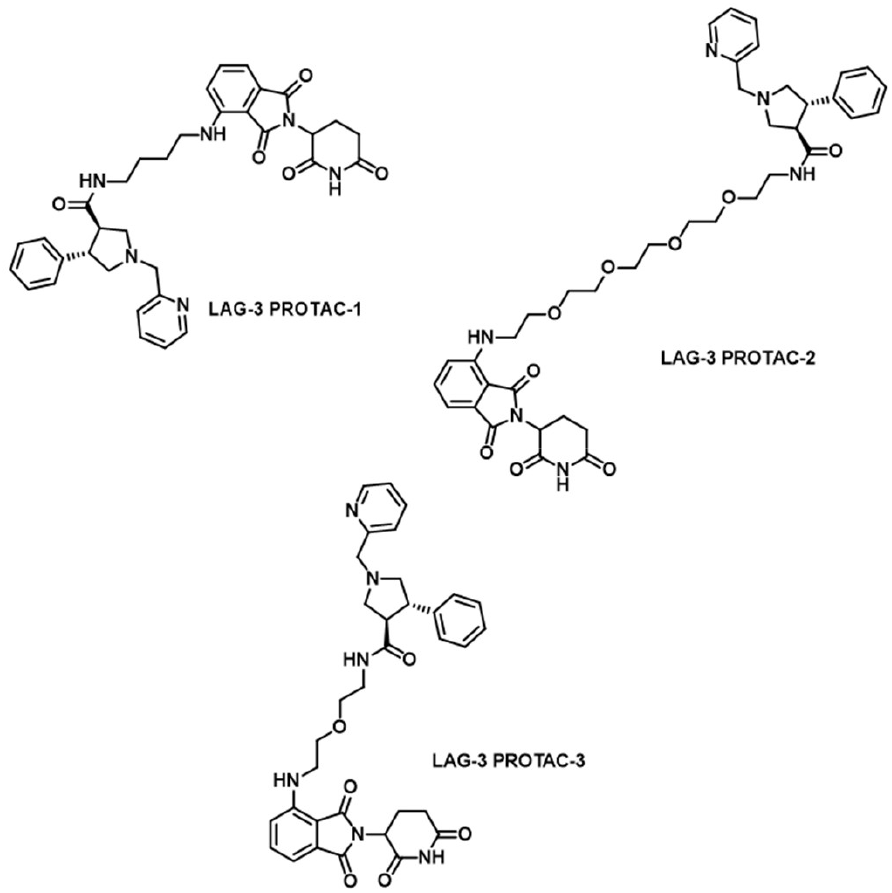
Chemical structures of CRL4^CRBN^-based PROTACs (**LAG-3 PROTAC-1, LAG-3 PROTAC-2, LAG-3 PROTAC-3**) employing pyrrolidine-3-carboxamide core scaffold and varying linker lengths.

We next evaluated the degradation activity of each PROTAC in Raji-LAG3 cells using Western blotting after 24-hour compound treatment (Figures 4A-C). **LAG-3 PROTAC-1** and **LAG-3 PROTAC-3** exhibited strong dose-dependent reduction of LAG-3 protein levels, while **LAG-3 PROTAC-2** showed negligible activity across all tested concentrations. These differences were not observed in Raji-Null cells lacking LAG-3, confirming target specificity. The lack of degradation by **LAG-3 PROTAC-2**, despite structural similarity to active compounds, highlights the critical role of linker optimization for successful E3 ligase recruitment and ternary complex stabilization.

**Figure 4.**
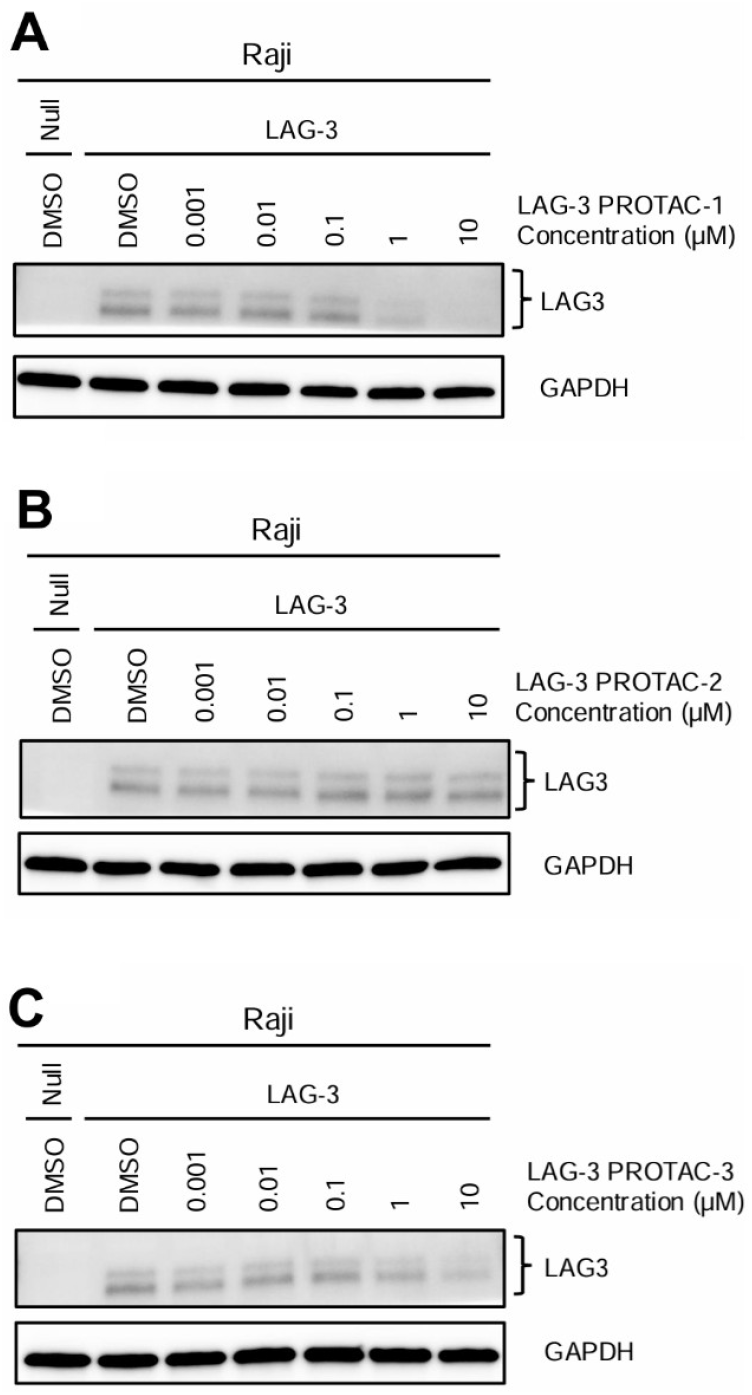
Western blot evaluation of LAG-3 degradation in Raji-LAG3 cells after 24 h treatment with **LAG-3 PROTAC-1 (A), LAG-3 PROTAC-2 (B)**, and **LAG-3 PROTAC-3 (C)**.

Densitometric analysis of Western blot bands from three independent experiments (Figure 5) confirmed the superior degradation efficiency of **LAG-3 PROTAC-1**, which reduced LAG-3 levels by up to 85% at 10 µM. **LAG-3 PROTAC-3** also induced significant degradation, though to a slightly lesser extent. **LAG-3 PROTAC-2** treatment resulted in minimal change in LAG-3 abundance, further reinforcing that linker length and positioning are key determinants of degrader potency. These differences are particularly interesting given that all three compounds share the same LAG-3-binding ligand and E3 ligase recruiter, indicating that linker composition alone can dramatically influence the formation and stability of the ternary complex. Such sensitivity to linker design is consistent with previous observations across diverse PROTAC systems, underscoring the context-specific nature of productive protein–ligase engagement. These results establish preliminary structure–activity relationships (SAR) that can guide future optimization of LAG-3-directed PROTACs and serve as a foundation for iterative design strategies aimed at improving degradation potency, selectivity, and drug-like properties.

**Figure 5.**
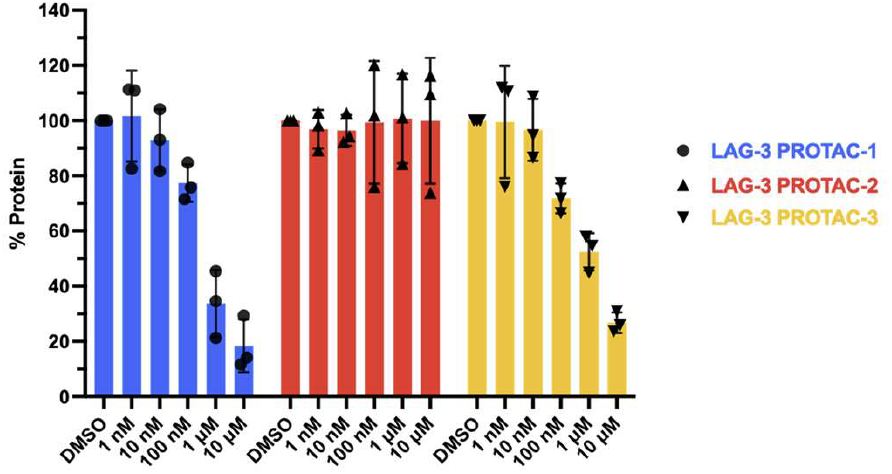
Quantification of LAG-3 western blot band signal after 24 h treatment with **LAG-3 PROTAC-1, LAG-3 PROTAC-2**, and **LAG-3 PROTAC-3** (n=3).

To quantify PROTAC potency, we determined the degradation half-maximal concentration (DC_50_) values after 24 hours of treatment (Figure 6). **LAG-3 PROTAC-1** demonstrated a DC_50_ of 274 nM, while **LAG-3 PROTAC-3** showed a slightly higher DC_50_ of 421 nM, consistent with their respective degradation profiles. These are within the typical range for first-generation LAG-3 PROTACs and suggest that our optimized ligand maintains sufficient affinity and spatial orientation to enable efficient ubiquitination of LAG-3. This represents the first quantitative demonstration of targeted degradation of an immune checkpoint protein using a small molecule PROTAC.

**Figure 6.**
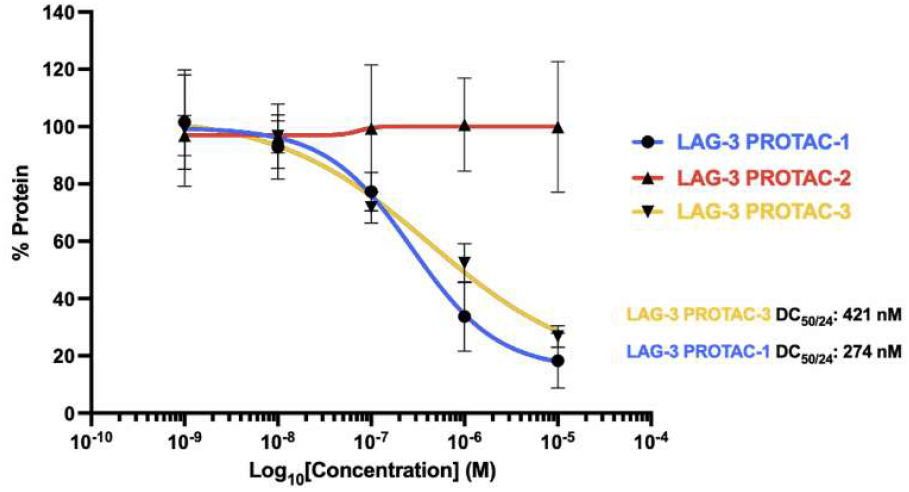
Half-maximal degradation concentration after 24 h (DC_50/24_) for **LAG-3 PROTAC-1** and **LAG-3 PROTAC-3**.

Given its superior degradation potency (DC_50_ = 274 nM) and robust activity profile across assays, **LAG-3 PROTAC-1** was selected as the representative compound for molecular docking and molecular dynamics (MD) simulations to investigate its LAG-3 binding mode. The surface view snapshots at 0, 50, and 100 ns (Figure 7 A-C) of **LAG-3 PROTAC-1** in complex with LAG-3 showed how **LAG-3 PROTAC-1** induces a conformational change in the LAG-3 binding pocket during the MD simulation. At the initial binding stage (0 ns) of the MD simulation, **LAG-3 PROTAC-1** adopts an extended conformation as it approaches the LAG-3 surface with minimal contact. However, as time goes by the warhead moiety anchors the PROTAC within the binding pocket, while the pomalidomide moiety remains positioned outside the pocket, poised to engage the E3 ligase (Figure 7A-E). The 50 ns snapshot (Figure 7B) captures an intermediate binding state where **LAG-3 PROTAC-1** has begun to establish key interactions with the LAG-3 binding site, showing partial insertion into the binding pocket with conformational rearrangement of both partners. By 100 ns, the complex reaches a stable, fully engaged configuration with optimal complementarity between the **LAG-3 PROTAC-1** warhead and the LAG-3 binding cavity and the pomalidomide moiety being situated in the solvent exposed region indicating its readiness to engage to the E3 Ligase.

**Figure 7.**
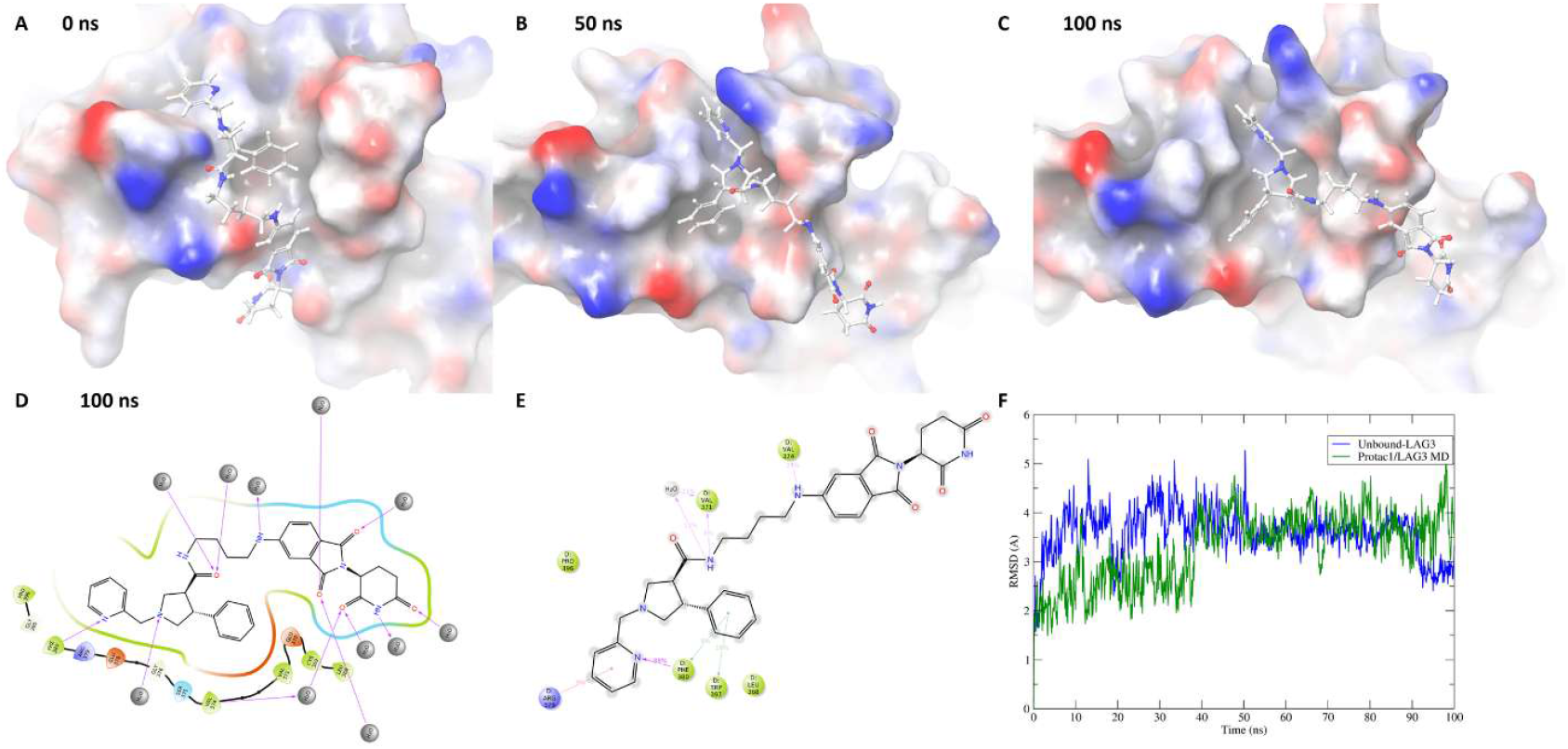
MD simulation analysis of LAG-3 PROTAC-1 binding to LAG-3. **(A-C)** Surface representations of LAG-3 (colored by electrostatic potential: blue = positive, red = negative, white = neutral) with PROTAC1 shown as stick representation at 0 ns **(A)**, 50 ns **(B)**, and 100 ns **(C). (D)** Two-dimensional interaction map of **LAG-3 PROTAC-1** with LAG-3 residues at 100 ns, showing hydrogen bonds (green lines) and hydrophobic contacts (pink and purple lines). **(E)** Contact frequency analysis displaying the percentage of simulation time that individual LAG-3 residues maintain contact with **LAG-3 PROTAC-1** throughout the 100 ns trajectory. **(F)** RMSD plots comparing structural stability of unbound LAG-3 (blue line) and **LAG-3 PROTAC-1**/LAG-3 complex (green line) over 100 ns simulation time. RMSD values are calculated relative to the initial structure after alignment of backbone atoms.

The 2D interaction diagram at 100 ns (Figure 7D) provides insights into the predicted interactions between the stabilized **LAG-3 PROTAC-1**/LAG-3 complex at 100ns. The analysis reveals a network of hydrogen bonds, hydrophobic contacts, and electrostatic interactions that anchor **LAG-3 PROTAC-1** within the LAG-3 binding site. Moreover, the hydrogen bond established between the warhead moiety and Phe380 amino acid residue of the binding pocket exhibited a stability of 88% throughout the 100ns MD simulation. The Root mean square deviation (RMSD) comparison (Figure 7F) demonstrates the structural stability of both the unbound LAG-3 protein (blue) and the **LAG-3 PROTAC-1**/LAG-3 complex (green). The unbound LAG-3 exhibits higher structural flexibility with RMSD values fluctuating between 3-5 Å with an average RMSD of 3.58 Å which is a reflection of the inherent flexible nature of LAG-3. In contrast, the **LAG-3 PROTAC-1**/LAG-3 complex shows reduced structural deviation with an average RMSD of 3.27 Å which is a reflection of the stabilizing effect of the **LAG-3 PROTAC-1** on LAG-3. The convergence of both trajectories after 60 ns suggests that the simulation has reached equilibrium, with the bound complex maintaining greater structural integrity than the unbound protein. This stabilization effect is characteristic of successful protein-drug interactions and supports the formation of a thermodynamically favorable complex.

This study presents the first report of small molecule PROTACs designed to target and degrade the immune checkpoint protein LAG-3. Starting from a DEL-derived LAG-3 binder, we synthesized a series of CRL4^CRBN^-based PROTACs and demonstrated that two compounds, **LAG-3 PROTAC-1** and **LAG-3 PROTAC-3**, effectively and selectively induce LAG-3 degradation in engineered cell models. The observed differences in degradation efficiency among the PROTACs underscore the critical influence of linker design on ternary complex formation and target engagement. Furthermore, molecular docking and MD simulations provided mechanistic insights into ligand binding and guided structure-based optimization. Together, these findings establish a foundational proof-of-concept for targeted degradation of LAG-3 and offer a new chemical strategy for modulating immune checkpoints. While these results demonstrate the feasibility of this approach, further optimization is required to improve PK properties and LAG-3 selectivity and to evaluate the functional consequences of LAG-3 degradation in primary immune cell models. This work lays the groundwork for future development of LAG-3 degraders with improved pharmacological properties and potential therapeutic application in cancer immunotherapy.

## Data Availability

Data will be made available on request.

## Acknowledgments

This work was supported by a grant from the ELSA U. Pardee Foundation (Award ID: 2022-215, PI: Gabr).

