## Supporting Information for "Design and Validation of the First-in-Class PROTACs for Targeted Degradation of the Immune Checkpoint LAG-3"

*Electronic Supplementary Information*

| **Contents** |  |
| --- | --- |
| Experimental  References | S1  S3 |
| Spectral data for **LAG-3 PROTAC-1-3** compounds | S4 |

**Experimental:**

**1. Chemistry:**

Commercially available chemicals and solvents were used without prior purification. The synthesized compounds were analyzed using ^1^H and ^13^C NMR, as well as high-resolution mass spectrometry (HRMS). ^1^H and ^13^C NMR spectra were acquired on a Bruker Avance III 500HD spectrometer (500 or 125 MHz, Billerica, MA, USA) using at room temperature. Chemical shifts are reported in parts per million (ppm) relative to tetramethylsilane (δ 0.00, s), with coupling constants given in hertz (Hz). Signal multiplicities are denoted as s (singlet), d (doublet), t (triplet), q (quartet), and m (multiplet). MS spectra were obtained using an SQ Detector 2 mass spectrometer (Waters, USA), while HRMS spectra were recorded on a Waters LCT Premier XE mass spectrometer (USA).

**General procedure for the synthesis of LAG-3 PROTAC-1-3.** To a stirred solution of **Fragment I** (1.0 equiv) in dry N,N-dimethylformamide (DMF) (0.1 M) was added HBTU (1.1 equiv) and triethylamine (3.0 equiv) at room temperature under a nitrogen atmosphere. The reaction mixture was stirred for 10 minutes to allow for activation of the carboxylic acid, after which the **Fragment II** (1.1 equiv) was added. The reaction mixture was stirred at room temperature for 12 hours, and progress was monitored by TLC. Subsequently, the reaction was quenched with water and extracted with ethyl acetate (3×50 mL). The combined organic layers were washed with brine, dried over anhydrous Na₂SO₄, filtered, and concentrated under reduced pressure. The crude product was purified by flash column chromatography (silica gel, gradient elution with ethyl acetate/hexanes) to afford the desired amide derivative.

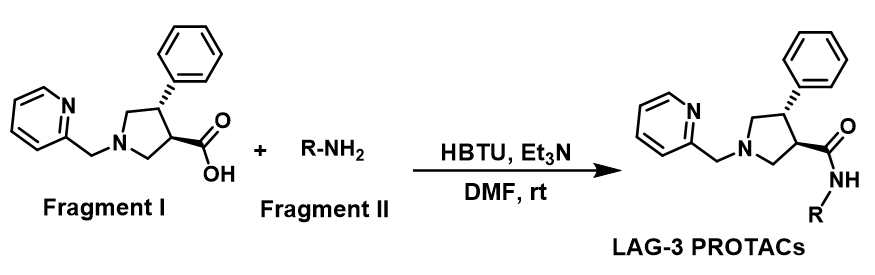

**Scheme 1.** Synthesis of the designed LAG-3 PROTACs based on amide coupling of Fragments **I** and **II**.

**LAG-3 PROTAC-1**. Yield 54%. (3*R*,4*S*)-*N*-(4-((2-(2,6-dioxopiperidin-3-yl)-1,3-dioxoisoindolin-4-yl)amino)butyl)-4-phenyl-1-(pyridin-2-ylmethyl)pyrrolidine-3-carboxamide. ^1^H NMR (500 MHz, CDCl_3_) δ ppm: 0.67– 1.01 (m, 1H), 1.20 (s, 2H), 1.50 – 1.64 (m, 1H), 1.98 – 2.11 (m, 1H), 2.60 – 3.34 (m, 13H), 3.53 (qd, *J* = 8.2, 3.8 Hz, 1H), 3.85 (dd, *J* = 16.3, 13.6 Hz, 1H), 4.02 (dd, *J* = 13.6, 3.4 Hz, 1H), 4.77 – 4.91 (m, 1H), 6.03 – 6.27 (m, 2H), 6.74 – 6.82 (m, 1H), 7.01 (d, *J* = 7.2 Hz, 1H), 7.11 – 7.15 (m, 2H), 7.17 – 7.23 (m, 4H), 7.31 – 7.45 (m, 2H), 7.61 (tt, *J* = 7.6, 1.4 Hz, 1H), 8.50 (d, *J* = 3.1 Hz, 1H), 8.94 (s, 1H). ^13^C NMR (126 MHz, CDCl_3_) δ ppm: 22.82, 23.30, 26.33, 26.88, 31.46, 38.76, 42.09, 48.50, 48.91, 53.31, 57.42, 60.68, 61.39, 109.98, 111.55, 116.69, 122.67, 123.34, 127.02, 127.50, 128.80, 132.49, 136.20, 136.92, 146.88, 149.32, 167.61, 168.61, 168.76, 168.83, 169.57, 170.27, 171.22, 171.35, 173.46, 173.53. HRMS (ESI): calculated for C_34_H_37_N_6_O_5_ (M + H)^+^, 609.2825; found, 609.2829.

**LAG-3 PROTAC-2**. Yield 63%. (3*R*,4*S*)-*N*-(14-((2-(2,6-dioxopiperidin-3-yl)-1,3-dioxoisoindolin-4-yl)amino)-3,6,9,12-tetraoxatetradecyl)-4-phenyl-1-(pyridin-2-ylmethyl)pyrrolidine-3-carboxamide. ^1^H NMR (500 MHz, CDCl_3_) δ ppm: 0.71 – 0.88 (m, 1H), 2.00 – 2.10 (m, 1H), 2.58 – 2.84 (m, 4H), 3.17 – 3.33 (m, 5H), 3.33 – 3.41 (m, 7H), 3.42 (d, *J* = 2.9 Hz, 2H), 3.43 – 3.47 (m, 1H), 3.50 (q, *J* = 2.7 Hz, 2H), 3.52 – 3.56 (m, 4H), 3.57 – 3.60 (m, 2H), 3.62 (q, *J* = 5.1 Hz, 3H), 3.65 – 3.75 (m, 1H), 4.17 (t, *J* = 12.6 Hz, 1H), 4.28 (t, *J* = 12.3 Hz, 1H), 4.79 – 4.90 (m, 1H), 6.43 (t, *J* = 5.6 Hz, 1H), 6.63 (d, *J* = 18.6 Hz, 1H), 6.82 (dd, *J* = 8.5, 1.9 Hz, 1H), 7.03 (d, *J* = 7.2 Hz, 1H), 7.11 – 7.18 (m, 2H), 7.21 – 7.32 (m, 5H), 7.38 – 7.43 (m, 1H), 7.51 – 7.65 (m, 3H), 8.41 – 8.54 (m, 1H). ^13^C NMR (126 MHz, CDCl_3_) δ ppm: 22.85, 29.71, 31.44, 39.41, 42.38, 47.73, 48.91, 52.52, 57.21, 59.58, 60.01, 69.36, 69.66, 70.25, 70.35, 70.49, 70.52, 70.61, 70.81, 110.32, 110.66, 111.65, 116.78, 118.59, 124.46, 125.72, 127.55, 127.74, 128.41, 128.95, 132.56, 136.04, 137.49, 146.84, 149.41, 167.64, 168.76, 168.88, 169.28, 171.48. HRMS (ESI): calculated for C_40_H_49_N_6_O_9_ (M + H)^+^, 757.3561; found, 757.3538.

**LAG-3 PROTAC-3**. Yield 58%. (3*R*,4*S*)-*N*-(2-(2-((2-(2,6-dioxopiperidin-3-yl)-1,3-dioxoisoindolin-4-yl)amino)ethoxy)ethyl)-4-phenyl-1-(pyridin-2-ylmethyl)pyrrolidine-3-carboxamide. ^1^H NMR (500 MHz, CDCl_3_) δ ppm: 2.59 – 3.74 (m, 19H), 4.17 (dd, *J* = 82.7, 13.3 Hz, 1H), 4.33 – 4.55 (m, 1H), 6.71 – 6.82 (m, 2H), 4.80 – 4.99 (m, 1H), 6.98– 7.19 (m, 7H), 7.23 – 7.61 (m, 4H), 7.69 (d, *J* = 8.1 Hz, 1H), 8.43 (d, *J* = 5.2 Hz, 1H). ^13^C NMR (126 MHz, CDCl3) δ ppm: 14.13, 22.83, 23.11, 29.51, 31.55, 39.54, 39.87, 41.14, 41.80, 49.15, 51.64, 57.46, 68.60, 109.65, 110.25, 110.94, 111.66, 116.78, 118.46, 124.26, 125.19, 127.11, 127.47, 127.63, 128.65, 132.42, 136.18, 136.51, 137.40, 137.89, 146.70, 149.20, 162.65. HRMS (ESI): calculated for C_34_H_37_N_6_O_6_ (M + H)^+^, 625.2775; found, 625.2765.

**2. Cell Culture for the Evaluation of LAG-3 PROTACs**. RAJI-Null and RAJI-hLAG-3 cell lines were purchased from Invivogen and cultured in RPMI 1640 medium. All media were supplemented with 10% fetal bovine serum (FBS) and 1% penicillin-streptomycin, and cells were incubated at 37°C in a humidified atmosphere containing 5% CO_2_.

**3. Western Blotting and Antibodies**. Protein extracts were prepared using RIPA buffer supplemented with protease inhibitors (Pierce Protease Inhibitor Mini Tablets, Thermo Fisher Scientific) and phosphatase inhibitors (Pierce Phosphatase Inhibitor Mini Tablets, Thermo Fisher Scientific). The insoluble fraction was removed by centrifugation (20,000g) for 10 min at 4°C. Protein concentrations in cell lysates were normalized using the Pierce BCA Protein Assay Kit (Thermo Fisher Scientific), following the manufacturer’s instructions. Proteins were separated by SDS-PAGE and transferred to PVDF membranes. Membranes were blocked with EveryBlot Blocking Buffer (Bio-Rad) and incubated with primary antibodies overnight at 4°C, followed by one-hour incubation of HRP-conjugated secondary antibodies. Enhanced chemiluminescence (SuperSignal West Pico Maximum Sensitivity Substrate, Thermo Fisher Scientific) was used for detection. The following antibodies were used: LAG-3 (1:1000, Proteintech Cat. No.: 16616-1AP), GAPDH (1:10,000. Proteintech Cat. No.: 60004-1-Ig)

**4. Molecular Docking.** The crystal structure of human LAG-3 domains 3-4 in complex with antibody single chain-variable fragment (PDB ID: 7TZH) was obtained from the protein data bank^1^. Induced Fit Docking (IFD) was performed using Schrödinger Suite to predict the binding pose of PROTAC1 with LAG-3. Chain D of LAG-3 was isolated and prepared using the Protein Preparation Wizard. The protein preparation step included optimization of hydrogen bond networks, assignment of protonation states at physiological pH, and energy minimization using the OPLS4 force field^2^. **LAG-3 PROTAC1** was prepared using the minimization module of Schrodinger to generate the most energetically stable conformation. The induced fit docking protocol was employed to account for receptor flexibility during ligand binding, allowing side chain movements within 5 Å of the ligand and backbone refinement of residues within 3 Å.

**5. Molecular Dynamics Simulation.** Molecular dynamics simulations were conducted using Schrödinger's Desmond MD system with the OPLS4 force field. The **LAG-3 PROTAC1**-LAG-3 complex was solvated in a TIP3P water model within an orthorhombic simulation box, maintaining a minimum distance of 10 Å between the protein surface and box boundaries. The system was neutralized with appropriate counterions (Na+ and Cl-) to achieve physiological ionic strength (0.15 M NaCl). The simulation protocol included an initial energy minimization followed by a series of equilibration steps: NVT equilibration at 300 K for 100 ps, followed by NPT equilibration at 300 K and 1 bar for 1 ns. Production runs were performed for 100 ns under NPT conditions using the Nosé-Hoover thermostat and Martyna-Tobias-Klein barostat. Trajectory frames were saved every 10 ps for analysis. An independent simulation of unbound LAG-3 was performed under identical conditions for comparison.

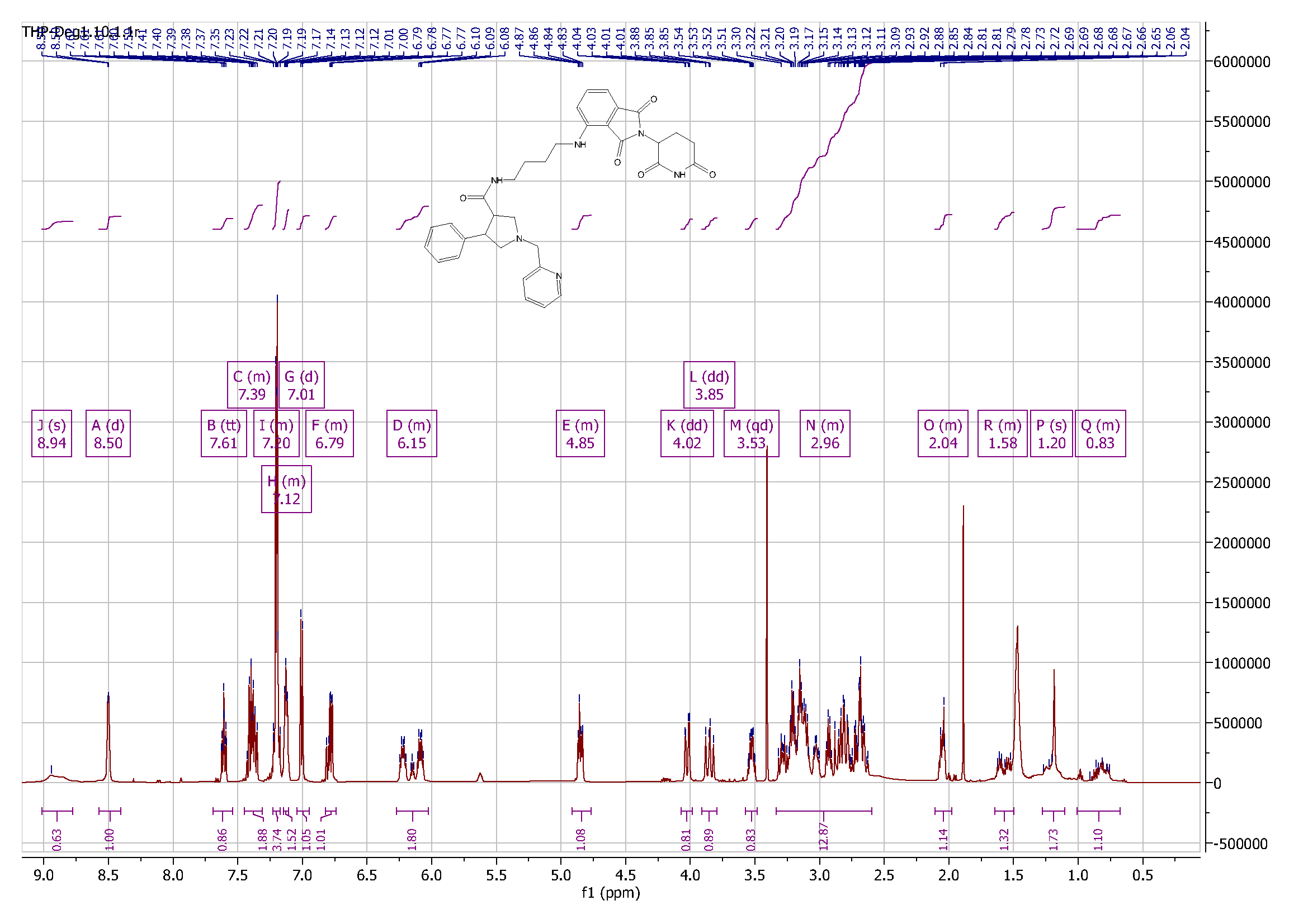

**Figure S1**. ^1^H NMR spectrum of **LAG-3 PROTAC-1**.

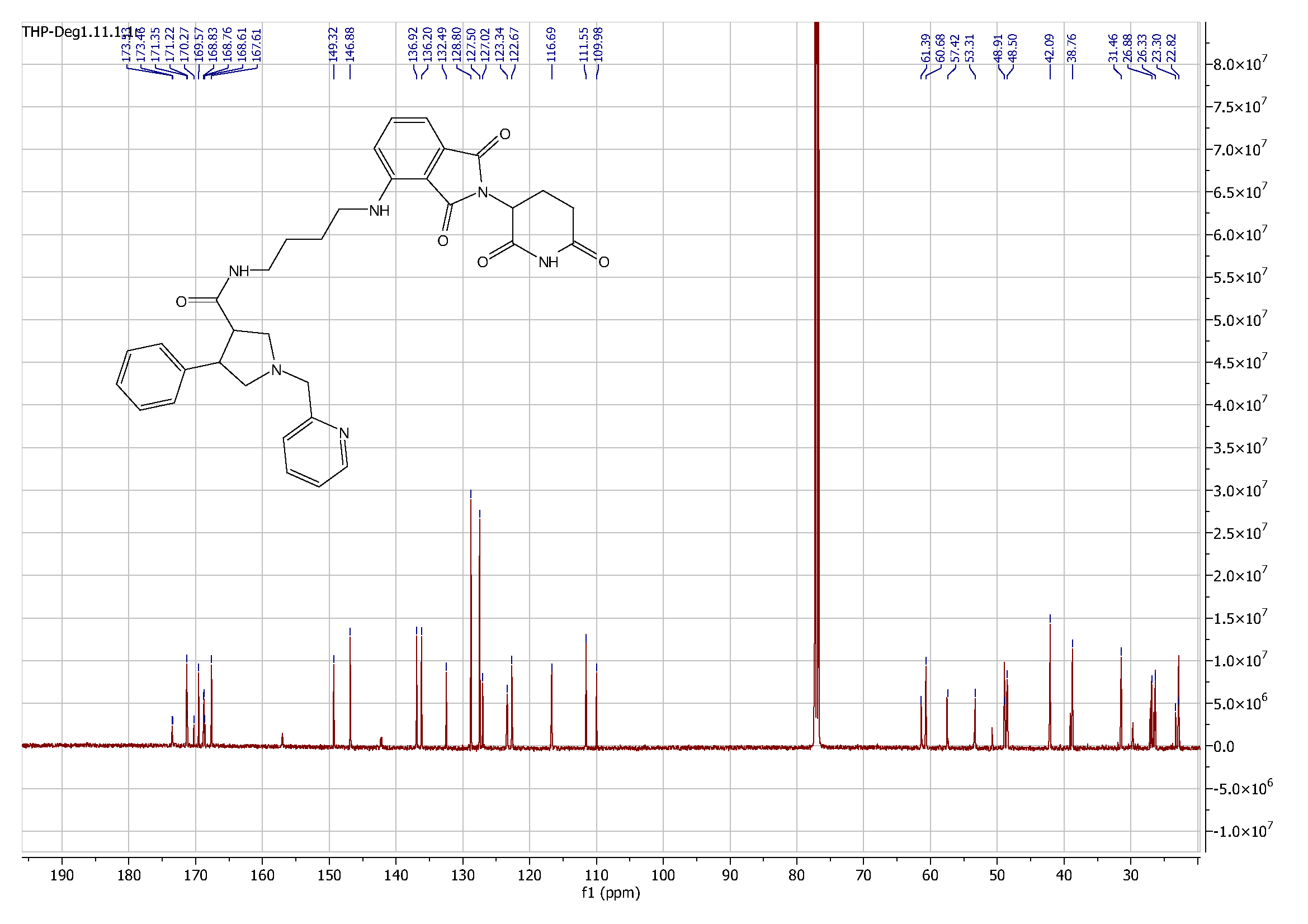

**Figure S2**. ^13^C NMR spectrum of **LAG-3 PROTAC-1**.

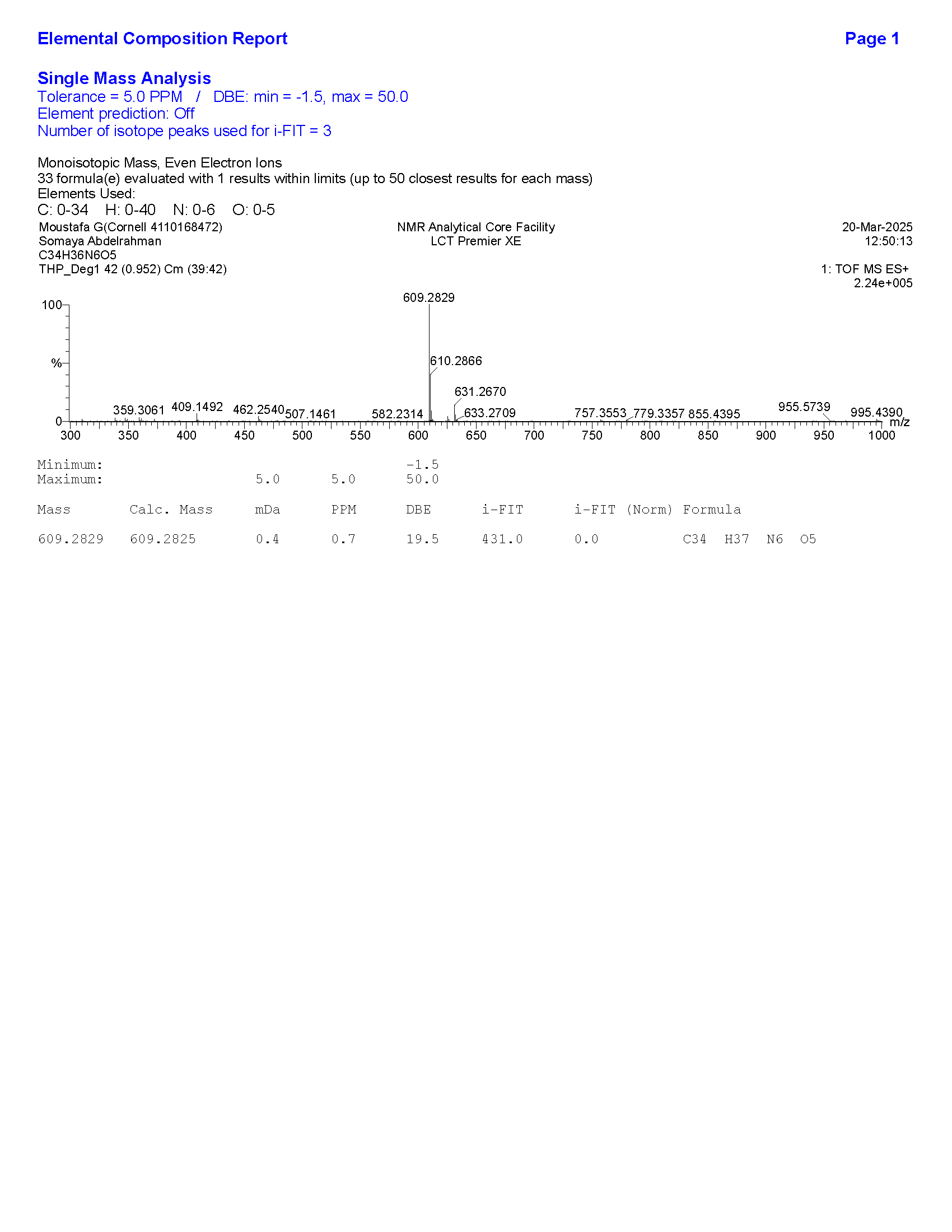

**Figure S3**. HRMS data of **LAG-3 PROTAC-1**.

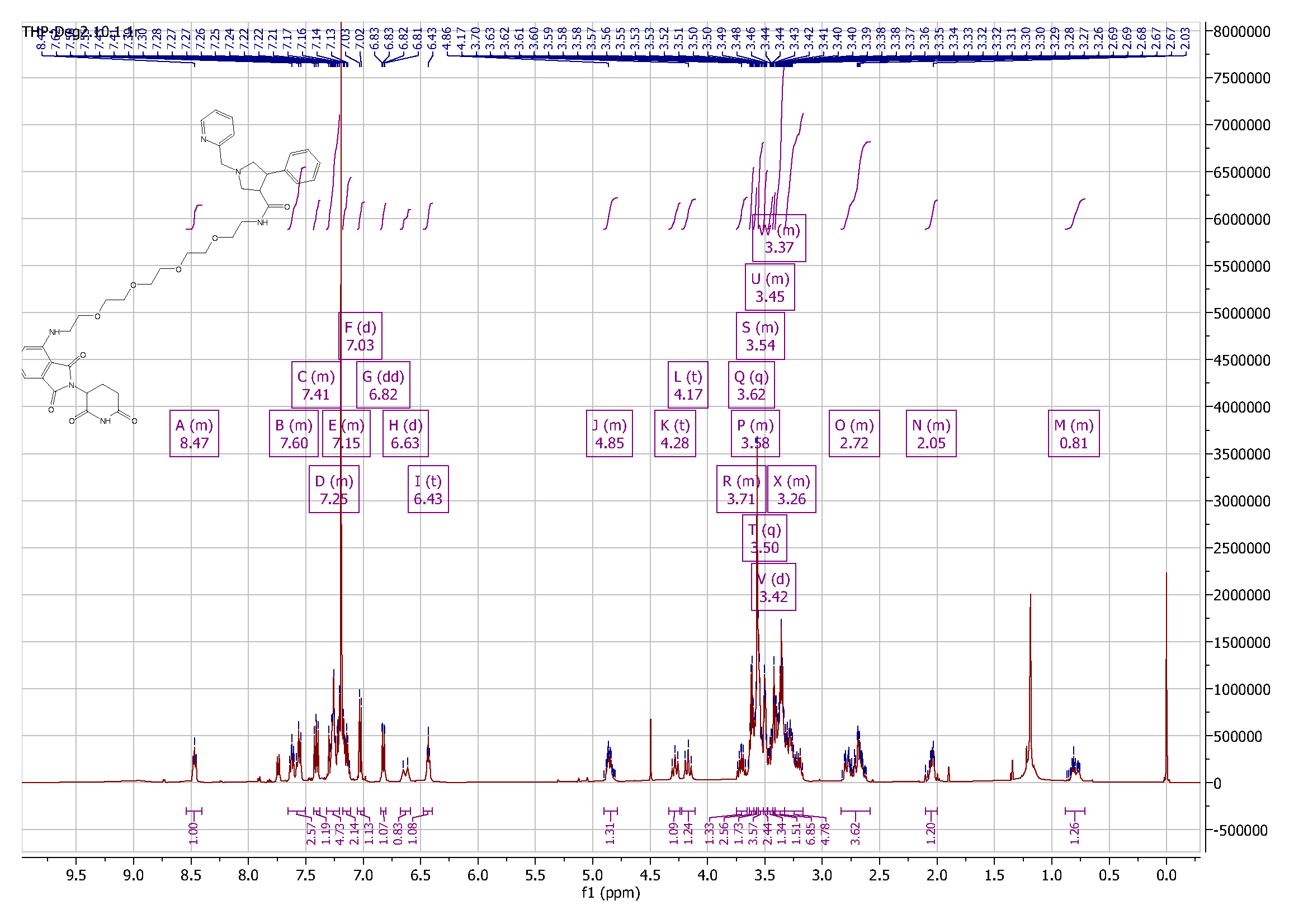

**Figure S4**. ^1^H NMR spectrum of **LAG-3 PROTAC-2**.

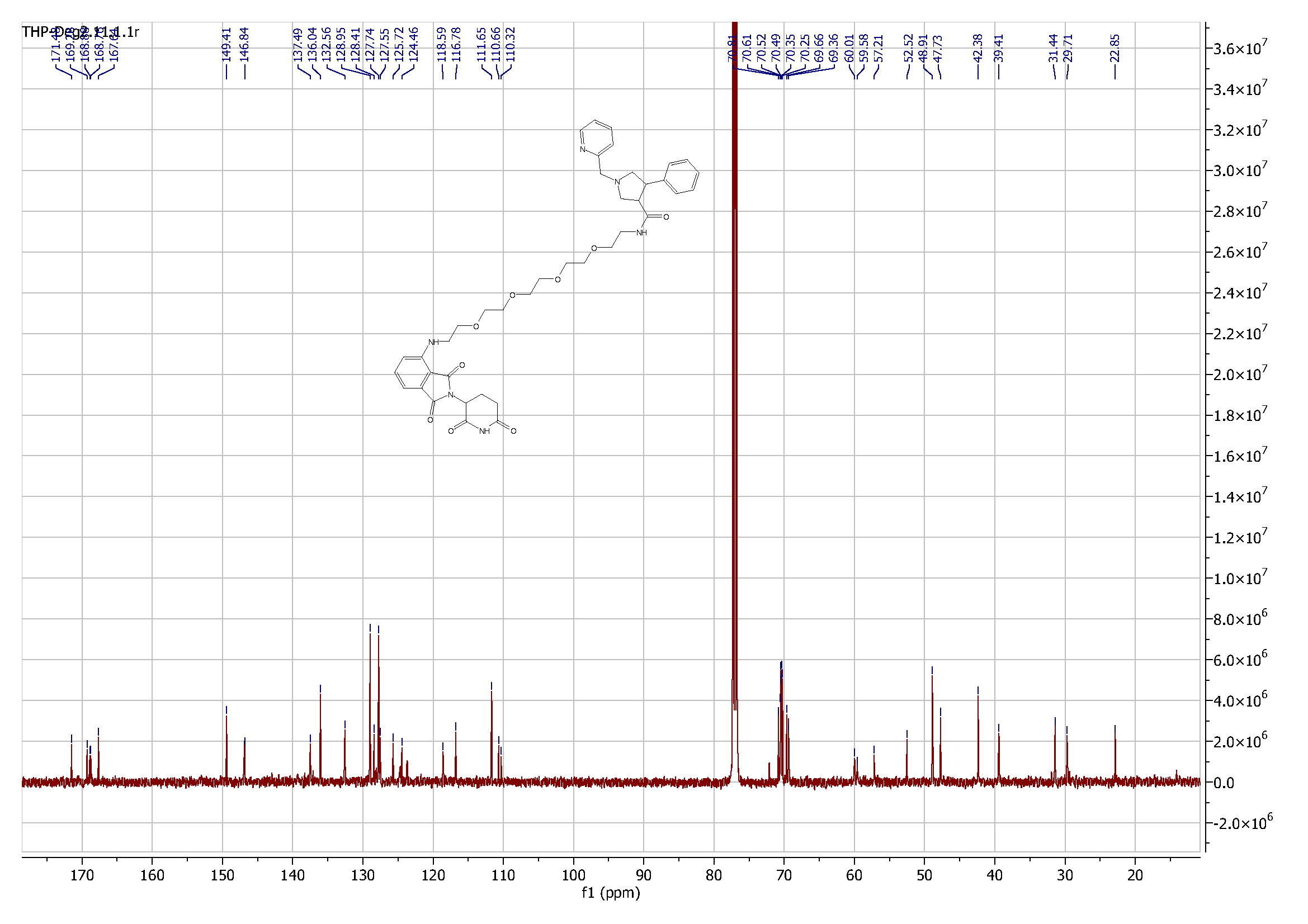

**Figure S5**. ^13^C NMR spectrum of **LAG-3 PROTAC-2**.

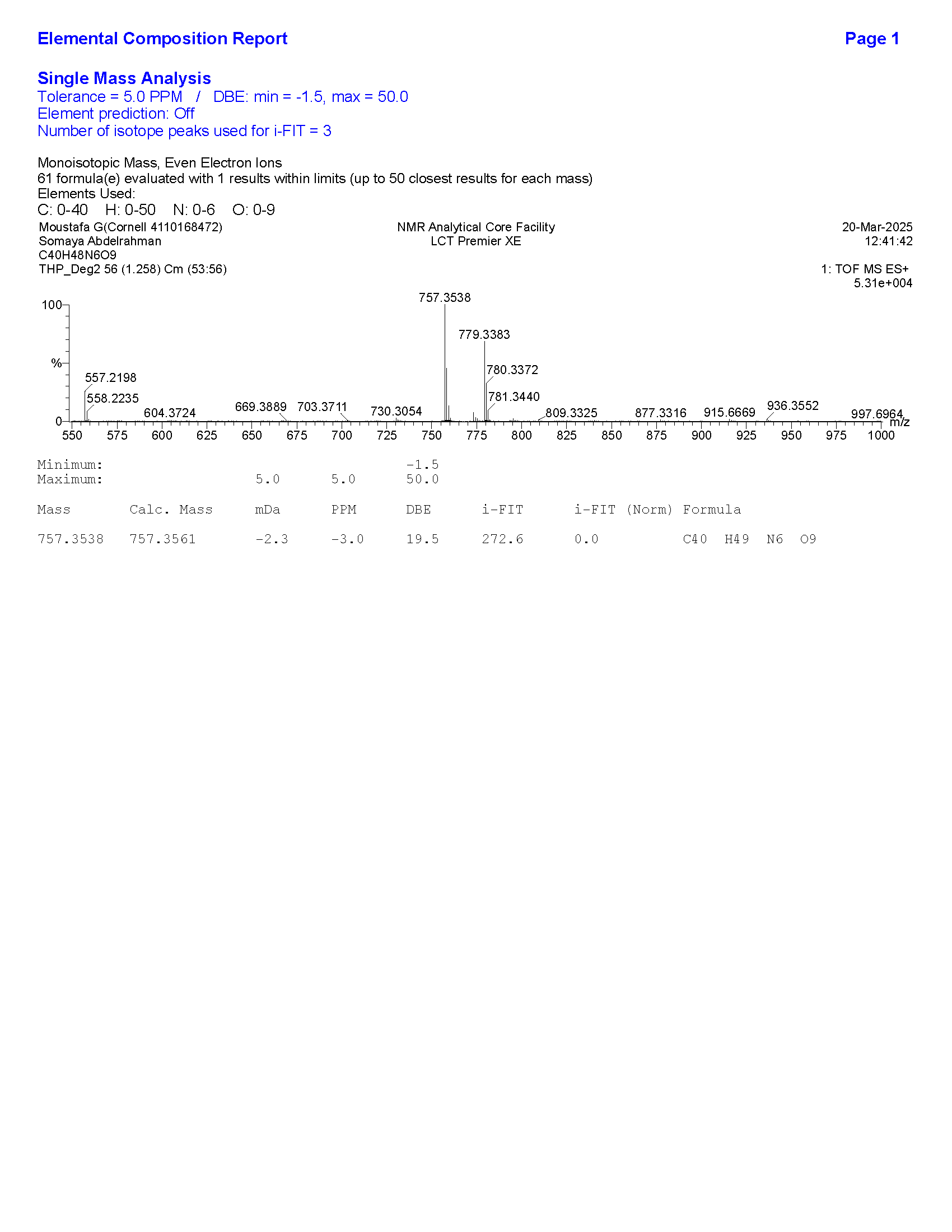

**Figure S6**. HRMS data of **LAG-3 PROTAC-2**.

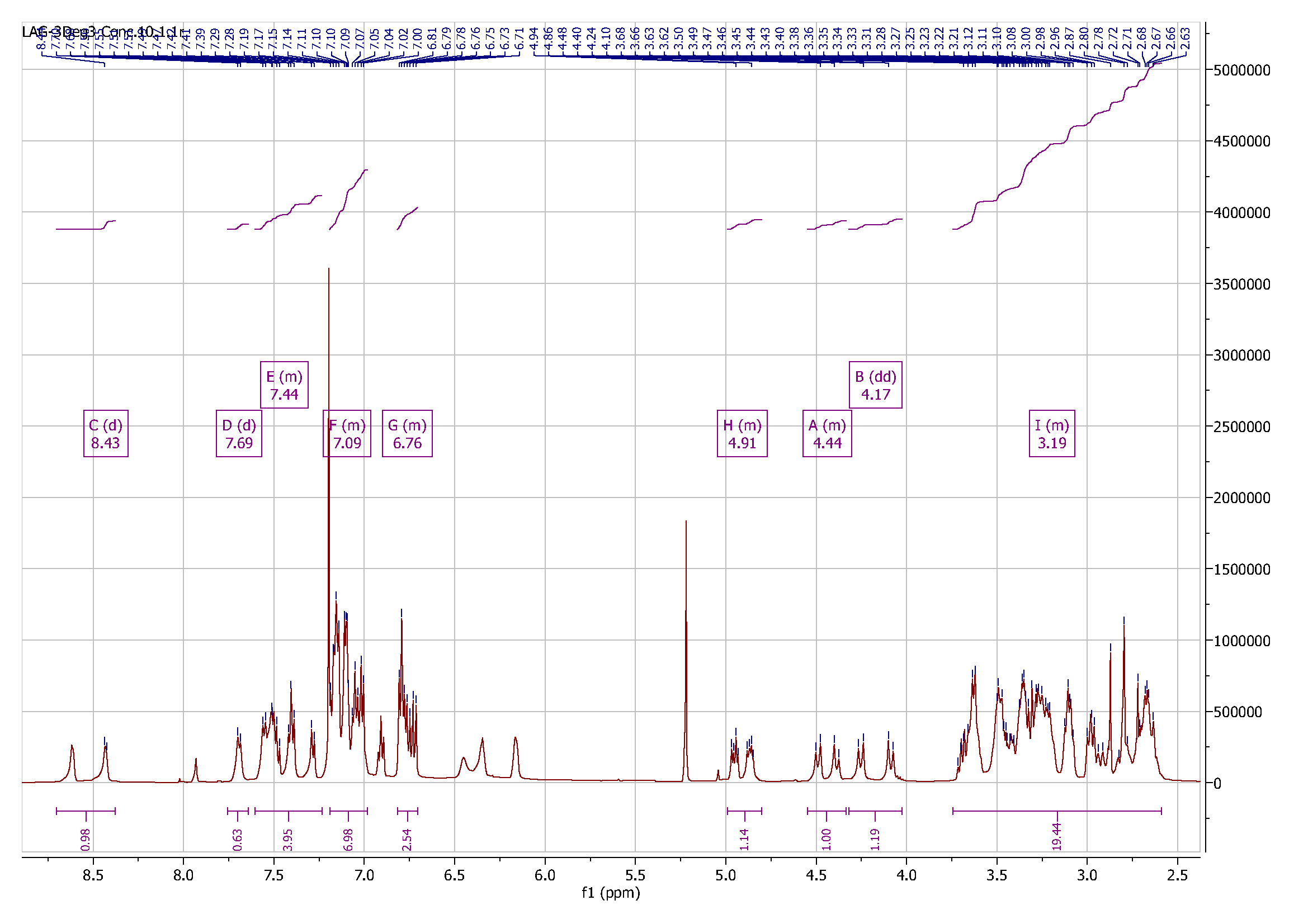

**Figure S7**. ^1^H NMR spectrum of **LAG-3 PROTAC-3**.

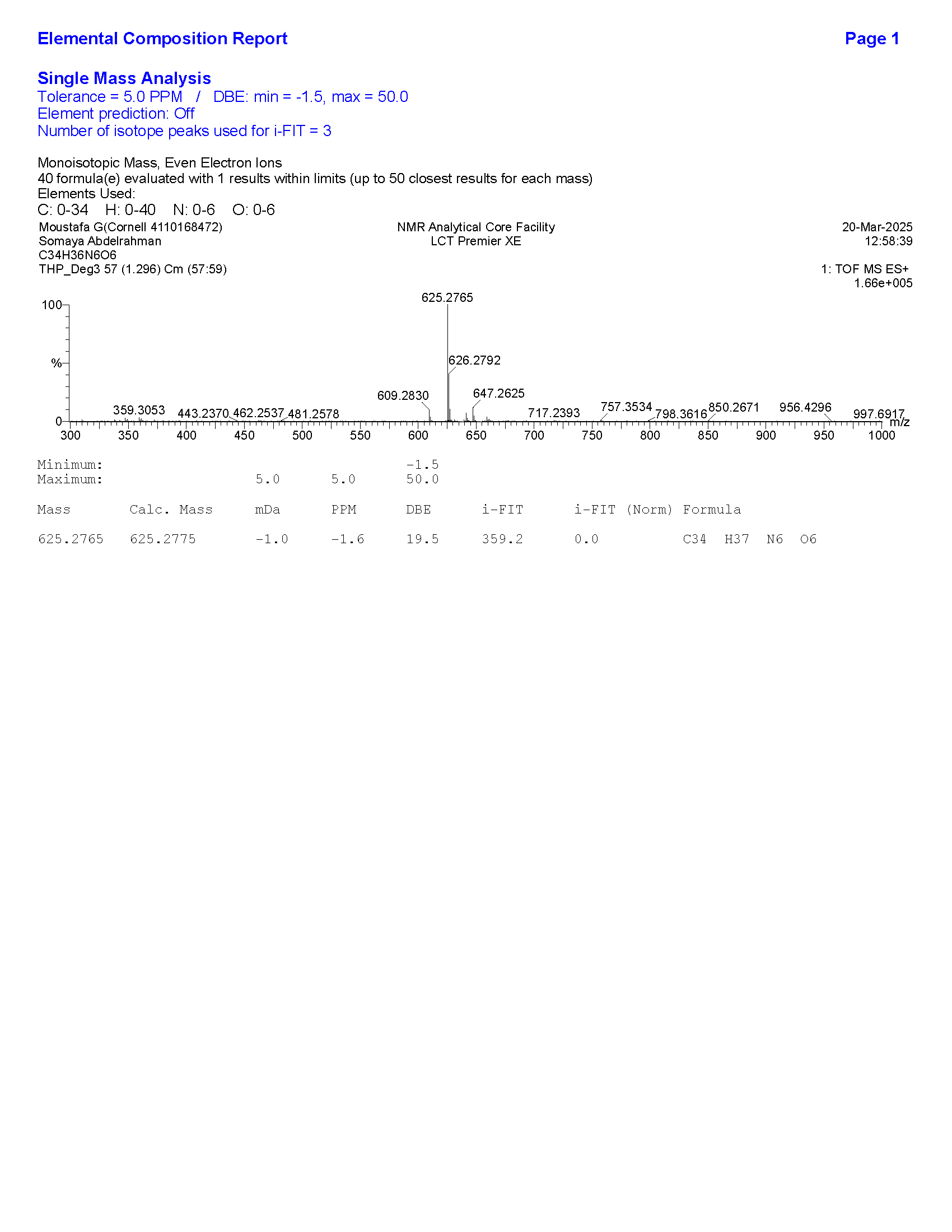

**Figure S8**. HRMS data of **LAG-3 PROTAC-3**.
